# Inferring disease-associated microRNAs using semi-supervised multi-label graph convolutional networks

**DOI:** 10.1101/666719

**Authors:** Xiaoyong Pan, Hong-Bin Shen

**Affiliations:** Institute of Image Processing and Pattern Recognition, Shanghai Jiao Tong University, and Key Laboratory of System Control and Information Processing, Ministry of Education of China, 200240 Shanghai, China; Department of Medical informatics, Erasmus Medical Center, 3015 CE Rotterdam, The Netherlands

## Abstract

MicroRNAs (miRNAs) play crucial roles in many biological processes involved in diseases. The associations between diseases and protein coding genes (PCGs) have been well investigated, and further the miRNAs interact with PCGs to trigger them to be functional. Thus, it is imperative to computationally infer disease-miRNA associations under the context of interaction networks.

In this study, we present a computational method, DimiG, to infer miRNA-associated diseases using semi-supervised Graph Convolutional Network model (GCN). DimiG is a multi-label framework to integrate PCG-PCG interactions, PCG-miRNA interactions, PCG-disease associations and tissue expression profiles. DimiG is trained on disease-PCG associations and a graph constructed from interaction networks of PCG-PCG and miRNA-PCG using semi-supervised GCN, which is further used to score associations between diseases and miRNAs. We evaluate DimiG on a benchmark set collected from verified disease-miRNA associations. Our results demonstrate that the new DimiG yields promising performance and outperforms the best published baseline method not trained on disease-miRNA associations by 11% and is also superior to two state-of-the-art supervised methods trained on disease-miRNA associations. Three case studies of prostate cancer, lung cancer and Inflammatory bowel disease further demonstrate the efficacy of DimiG, where the top miRNAs predicted by DimiG for them are supported by literature or databases.

## 1 Introduction

MicroRNAs (miRNAs) are a type of small non-coding RNAs with size about 22 nucleotides, which interact with other RNAs to play important roles in transcriptional and post-transcriptional gene regulation (Bartel, 2004). It is estimated that over 60% of all human protein coding Genes (PCGs) are regulated by miRNAs (Friedman, et al., 2009), and these miRNAs have been implicated in diseases. To date, the associations between diseases and PCGs are well investigated, many disease-PCG associations have been discovered and collected in public databases, e.g. DISEASES (Pletscher-Frankild, et al., 2015), OUGene (Pan and Shen, 2016) and DisGeNET (Pinero, et al., 2017). Compared to PCG’s well known important roles in diseases, the studies of effects of miRNAs are still few. With increasing high-throughput sequencing data generated, more and more miRNAs are being discovered, experimentally identifying their functions is costly and time-consuming. Thus, it is imperative to develop computational methods to identify functional miRNA biomarkers associated with diseases, especially using rich information buried in PCGs.

Some miRNAs are mainly expressed in certain tissues and show tissue specificity (Ludwig, et al., 2016), which have certain tissue-specific expression patterns associated with diseases (Baker, et al., 2017). They are expected to behave similarly to other disease-associated genes like PCGs or long non-coding RNAs (lncRNAs). Thus, several existing computational methods have used tissue expression data to infer gene-disease associations. For instance, the GeneTIER makes use of disease-tissues associations to prioritize disease candidate genes (Antanaviciute, et al., 2015); the NetWAS identifies disease-associated genes through combining tissue-specific interaction networks and genome-wide association study (GWAS) (Greene, et al., 2015). Also, some methods use tissue expression profiles with machine learning models to infer disease-associated lncRNAs. For example, the Dis-lncRF trains machine learning models on tissue expression profiles of disease-associated PCGs and further applies the trained models to infer disease-associated lncRNAs (Pan, et al., 2018). All the above studies demonstrated that tissue expression profiles indeed can facilitate the detection of disease-gene associations.

On the other hand, interaction networks contain rich clues for linking miRNAs to diseases. Many computational methods have been developed under the context of gene-gene networks (Chen, et al., 2017). For example, Jiang el al. integrate miRNA and disease similarity network, and miRNA-disease association to prioritize disease candidate miRNAs using a network-based approach (Jiang, et al., 2010); the RWRMDA implements random walk on the miRNA functional similarity network to link miRNAs to diseases (Chen, et al., 2012); the MDHGI integrates the predicted association score based on sparse learning method to infer disease-associated miRNAs (Chen, et al., 2018). More closely related studies include: DRMDA applies stacked autoencoder to learn deep representation for predicting miRNA-disease association (Chen, et al., 2018); LRSSLMDA and PBMDA uses Laplacian regularized sparse subspace learning and path-based computational model for miRNA-disease association prediction (Chen and Huang, 2017; You, et al., 2017); BNPMDA uses Bipartite Network Projection based on the known miRNA-disease associations (Chen, et al., 2018); the KBMF-MDI employs kernelized Bayesian matrix factorization to score miRNA-disease associations by integrating disease and miRNA similarity (Lan, et al., 2018). Similarly, DLRMC infers disease-associated miRNAs using Dual Laplacian regularized matrix completion (Tang, et al., 2019).

A common hypothesis for the above methods is they assume that similar miRNAs can be associated with the same disease and similar diseases would be associated with the same miRNA. Thus, they commonly train and evaluate the models with representations of miRNAs and diseases as inputs on verified disease-miRNA associations through cross-validation approach.

However, as pointed out in (Lehtinen, et al., 2015), in the context of gene function prediction, cross-validation may be problematic since some gene-function associations are not independent in the benchmark set. There exists the same issue for disease-miRNA associations due to the following: 1) MiRNAs from the same family maybe associated with the same disease; 2) Disease-associated miRNAs from miRNA-target assay maybe derived from the targets these miRNAs interact; 3) The associated miRNAs of child diseases are also related to the miRNAs of parent diseases in disease ontology. When training and evaluating the models using cross-validation, randomly dividing the disease-miRNA associations may cause dependent associations separately into the training and test set, potentially leading to an overestimated predictive performance. Cross-validating miRNA-disease associations may not actually reflect the method’s ability to predict new miRNA-disease associations, but rather which information is dissipated in the benchmark set. In addition, as reported in (Park and Marcotte, 2012), there may exist flaws in cross-validation for computational pair-input prediction. One disease or one miRNA may be associated with multiple miRNAs or diseases, so randomly dividing disease-miRNA pairs into training and test set will make some pairs in the test set share either the miRNA or the disease with the pairs in the training set, which causes the trained models not generalize well to unseen disease-miRNA associations.

Thus, during cross-validation, complicate steps are required to make sure that dependent samples are divided into the same training set or the same test set and pairs in the training and test set do not share the miRNA or disease. It is almost impossible to construct a completely independent test set. An alternative strategy is that we do not use disease-miRNA associations for model training. For instance, instead of using miRNA-disease associations for model training, the miRPD approach combines PCG-disease associations and miRNA-PCG network to score miRNAs and diseases (Mork, et al., 2014). This has triggered us to further investigate disease-miRNA associations based on an interaction network. To date, there exists many high-confident disease-PCG associations, and one miRNA may share the same disease with its PCG targets, we will be capable of transferring PCG associated diseases to miRNAs on an interaction network under a new semi-supervised framework.

Recently, deep learning has achieved remarkable results in computational biology (Angermueller, et al., 2016; Ching, et al., 2018), especially convolutional neural networks (CNNs) (Lecun, et al., 1998). CNNs can capture local correlation buried in data and mainly consist of convolutional layers, pooling layers and fully connected layers. Tremendous studies have demonstrated the CNN networks are powerful in learning the hidden patterns from complicated biological data. For example, the DeepBind (Alipanahi, et al., 2015) and DeepSEA (Zhou and Troyanskaya, 2015) apply CNNs to predict preference of DNA/RNA-binding proteins and the impact of non-coding variants, respectively. The iDeep (Pan and Shen, 2017) and iDeepE (Pan and Shen, 2018) further improve the performance of predicting RNA-binding protein (RBP) binding sites and motifs using hybrid CNNs. The iDeepS (Pan, et al., 2018) identifies binding sequence and structure preferences of RBPs simultaneously using CNNs and long short temporary memory network (LSTM).

Although the CNN has shown its power, it cannot handle structured datasets, like gene-gene networks. To analyze these types of network data, graph convolutional networks (GCNs) have been developed (Defferrard, et al., 2016; Hamilton, et al., 2017; Kipf and Welling, 2017). Under the framework of spectral graph convolutions, it encodes both local graph structure and features of nodes. The GCNs have been used on the graph data to predict polypharmacy side effects, where the graph is a multimodal graph constructed from protein-protein interactions, drug-protein interactions, and the polypharmacy side effects (Zitnik, et al., 2018). The GCN is a graph-based semi-supervised learning method that does not require labels for all nodes. This setting is especially powerful for inferring miRNA-associated diseases, since many miRNAs are not well investigated about their associations with diseases. In addition, one PCG or miRNA can be associated with multiple diseases. Thus, we can formulate the prediction of disease-miRNA associations as a multi-label classification problem.

In this study, we present a new semi-supervised multi-label learning method, DimiG, based on GCNs to integrate multiple networks of PCG-PCG interactions, PCG-miRNA interactions, PCG-disease associations and tissue expression profiles to infer miRNA-associated diseases. The DimiG does not require the disease-miRNA associations, and it is trained on the graph consisting of PCG-PCG and miRNA-PCG interactions, where only PCGs have labelled diseases. Then DimiG is further used to score associations between diseases and miRNAs.

This study has made the following four major contributions for understanding disease-miRNA associations: 1) We further demonstrate that cross-validation performance of methods trained on known disease-miRNA associations could be overestimated and may not be able to reflect the method’s actual ability to predict new disease-miRNA associations. We have proposed a network-based knowledge transfer approach for this problem. Considering that a miRNA may share the same disease with its PCG targets and there exist many high-confident disease-PCG associations, we will be able to transfer the PCG associated diseases to miRNAs in an interaction network framework. 2) We have formulated disease-miRNA association prediction as a semi-supervised multi-label node classification in a graph, which can help learn the complex networks composed by unlabeled miRNAs and labeled PCGs and the multi-label associations. This is a new prediction protocol for this problem. 3) We use semi-supervised GCN to learn from PCG-associated diseases on an interaction network, which is further used to score diseases and miRNAs. This GCN-based approach combines the advantages of deep learning for representation learning and network-based methods. 4) We have further incorporated the domain knowledge into our model construction. Considering that miRNAs are often expressed in a tissue-specific way, we integrate the expression profiles across tissues into our GCN framework. Our results demonstrate that informative signals in more tissues can be captured for aiding the inference of disease-associated miRNAs.

## 2 Methods

We collect genome-wide tissue expression profiles from RNA-seq data, disease-associated PCGs, PCG-PCG interactions, miRNA-PCG interactions. These data are fed into a semi-supervised multi-label GCN using PCGs as training and validating. Then the trained GCN model is propagated into miRNAs to score their associations with diseases.

### 2.1 Data sources

One recent benchmark study (Huang, et al., 2018) shows that among 21 widely used protein-protein interaction databases, STRING (Szklarczyk, et al., 2015) is one of the three databases having the best performance for discovery of disease genes. Like the STRING database, miRNA-gene interaction database RAIN (Junge, et al., 2017) and disease-gene association database DISEASES (Pletscher-Frankild, et al., 2015) also integrate different sources of data, including text mining, knowledge, experiments and predictions using similar techniques and scoring schema, and they are developed by the same group and use the same gene identifiers. For disease-miRNA associations, the widely used HMDD database (Li, et al., 2014) is used as the benchmark set for evaluating disease-miRNA association prediction.

#### 2.1.1 Tissue expression data

We download the GTEx tissue expression data (GTEx_Analysis_v6p_RNA-seq_RNA-SeQCv1.1.8) across 53 tissues (Lonsdale, et al., 2013). It contains 19,732 PCGs and 2,833 miRNA genes. Of the 2,833 miRNAs, only 1,244 miRNAs are in miRBase v20 (Kozomara and Griffiths-Jones, 2011). In addition, we also collected three raw RNA-seq datasets, E-MTAB-513 across 16 tissues (Derrien, et al., 2012), GSE43520 across four tissues (Necsulea, et al., 2014) and GSE30352 across six tissues (Brawand, et al., 2011). The raw reads are mapped to the same reference genome as GTEx using STAR version 2.5.0b (Dobin, et al., 2013) and quantified using Cufflinks (Trapnell, et al., 2014). For a tissue with multiple samples, we calculate the median expression values for this tissue. The expression values are log-transformed using log_2_(1 + *x*), which is further normalized across tissues by the fraction of expression of one gene in one tissue relative to the sum of its expression in all tissues. In addition, we add another feature of the sum of one gene’s expression across all tissues.

#### 2.1.2 Disease-PCG associations

We download the integrated disease-PCG associations from DISEASES database (Pletscher-Frankild, et al., 2015). DISEASES database has multiple channel of evidences, e.g. knowledge, experiments and text mining, to support the associations. Each disease-PCG association is assigned a confidence score. In this study, we only use associations with confidence scores greater or equal to 2. All gene names are mapped to Ensembl gene identifiers and diseases are represented in Disease Ontology ID (Schriml, et al., 2012). In total, we obtain 86,430 associations between 4,161 diseases and 9,636 PCGs. Supplementary Fig. S1A shows the disease label distribution for PCGs.

#### 2.1.3 Disease-miRNA associations

We extract the disease-miRNA associations from database HMDD v3.0 (Li, et al., 2014), which has 32,281 associations between 1,102 miRNA genes and 850 diseases. After removing some associations whose disease name have no DOID, we build a set with 24,320 unique disease-miRNA associations between 1,007 miRNAs and 582 diseases. We further map the miRNA names into Ensembl gene identifiers. In the end, we obtain 6,829 associations between 548 miRNAs and 486 diseases. Supplementary Fig. S1B illustrates the disease label distribution for miRNAs.

#### 2.1.4 PCG-PCG interactions

The human gene-gene interactions are extracted from widely used database STRING v10 (Szklarczyk, et al., 2015), which houses millions of gene-gene interactions across multiple species. In STRING, each interaction is scored according to multiple evidences, including experiments, knowledge, prediction and text mining. The higher the score is, the more reliable this interaction is. We mapped all gene names to Ensembl gene ID and only select those interactions with confidence score greater than 400 (confidence values are between 1 and 1000), which is the medium confidence score in STRING database. In total, we obtain 1,481,757 interaction pairs of 18,883 PCGs.

#### 2.1.5 miRNA-PCG interactions

We extract the human miRNA-PCG interactions from RAIN database (Junge, et al., 2017), which integrates miRNA-gene interactions from text mining, experiments, knowledge and predictions. In total, RAIN scores 46,472 human miRNA-gene interactions according to different evidences. In this study, we only select those interactions with combined score greater than 0.15, which is used as default cut-off in the webserver. We mapped all miRNA names to Ensembl gene ID. In total, we obtained 173,662 interaction pairs between 17,686 PCGs and 1,725 miRNAs.

### 2.2 Data processing

We integrate the above tissue expression profiles, PCG-PCG interactions, miRNA-PCG interactions and PCG-disease associations to score diseases and miRNAs. When constructing the DimiG, we process the data as the following: 1) when constructing a graph, each PCG has at least one interacting PCG and each miRNA has at least one interacting PCG; 2) both the PCGs and miRNAs in the graph have expression profiles across tissues as node features; 3) the PCGs in the graph should be associated with at least one disease, since diseases are the labels of PCG nodes and they are used to calculate the training loss for model training. The whole processing is shown in Fig. 1. Finally, we obtained 7,222 genes with 6,188 PCGs and 1,034 miRNAs, and 248 diseases as the labels.

**Fig. 1.**
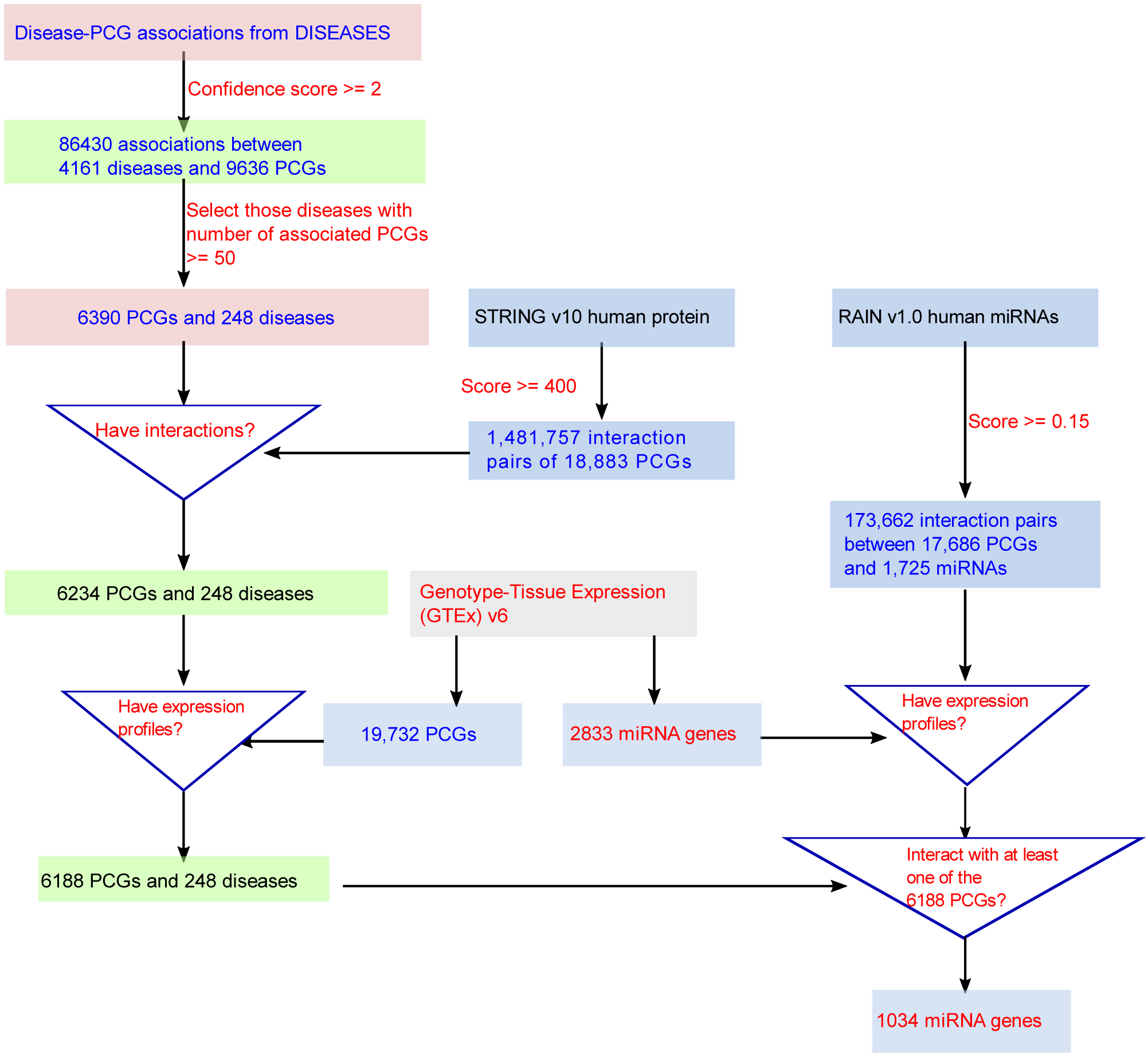
The flowchart of data processing for constructing the benchmark set for DimiG. All the gene names are mapped to Ensembl gene IDs and disease names are mapped to Disease Ontology ID.

Here, we use the experimentally verified miRNA-disease associations to evaluate the performance of DimiG. After removing those associations whose disease is not among the 248 diseases, we obtain 2,695 associations for 91 diseases. As negative control examples, for each disease, we randomly select the same number of miRNAs not associated with the disease in HMDD v3.0 as we have miRNAs associated with it. In the end, we obtain a dataset with 5,390 disease-miRNA pairs, where half of them are positives and the other half are negatives. This dataset is balanced not only overall but also for each disease, and is used as independent test set, whose association pairs are not involved in model training.

### 2.3 Graph Convolutional Networks

GCN (Kipf and Welling, 2017) is trained in an end-to-end way and learns the informative node embedding for the semi-supervised classification tasks on a graph. The GCN is different from previous network embedding methods, e.g. node2vec (Grover and Leskovec, 2016), which needs firstly to learn node embedding from a graph, and then the learned embedding is fed into a supervised classifier for downstream classification tasks.

Given a graph G and additional node feature vectors *X*:

1. A node feature matrix *X*: a *N* × *D* feature matrix, where *N* is the number of nodes in the graph G, and *D* is number of input features.
2. An adjacency matrix *A*: a *N* × *N* matrix, showing the connections between nodes in the graph G.
3. The multi-hot encoded label matrix Y for some nodes but not all nodes in the graph, one node can have multiple labels.

The GCN frames the classifying nodes in a graph as graph-based semi-supervised learning, where not all nodes in the graph need labels. To this end, a graph Laplacian regularization term is introduced into the loss function *L*:

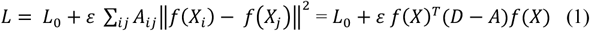

where *L*_0_ is the supervised loss, *f* can be a neural network-based function, *ε* is the factor, *X* is the node features, *A* is the adjacency matrix and *D* is the degree matrix.

A multi-layer GCN is propagated with the following rule:

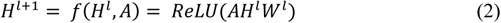

where *H*^*l*^ is the output of layer *l* and *W*^*l*^ is a weight matrix.

The GCN performs spectral convolutions on the graph and perform neighborhood weighting. A first-order approximation of localized spectral filters in Fourier-domain can be used to approximate the propagation rule (Defferrard, et al., 2016). Please refer to (Kipf and Welling, 2017) for more details on the GCN.

For semi-supervised classification, we minimize the binary cross entropy loss on labelled data:

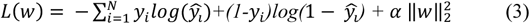

where y_i_is the true label, 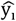 is the output probability of the last sigmoid layer and α is the weight factor for regularization.

### 2.4 Multi-label classification

Multi-label classification assigns multiple labels to one node, and it is categorized into two groups: (1) problem transformation methods and (2) algorithm adaptation methods. For problem transformation methods, they are transformed into multiple binary classification problems, where each binary classifier is trained separately for each label. For algorithm adaptation methods, they usually perform multi-label classification directly using one classifier, which outputs predicted probabilities for all the labels.

In DimiG’s multi-label assignment, we use the 2^nd^ approach and set up a sigmoid layer as the last output layer, whose number of neurons is the number of labels. Given a set of labels with *K* classes (248 diseases in this study), each node is assigned with several labels from it. Denote *x* as an input node, and *y* as a multi-hot encoded vector with 248 elements, which indicates absence of class *i* and presence of class *i*: *y* = [0, 1, 0, …, 1].

### 2.5 DimiG pipeline

DimiG integrates gene network, expression profiles and PCG-disease associations using semi-supervised multi-label GCN. DimiG only requires the PCGs with associated diseases as labels during the model training, then it propagates the node embedding to those miRNAs and further infer their associated labels. DimiG consists of two layers of GCNs, the node features are expression profiles of PCGs and miRNAs, the adjacency matrix is derived from PCG-PCG and PCG-miRNA interactions, and the labels are the associated diseases with PCGs. In the end, DimiG outputs a 1034 ×248 score matrix, where 1034 is the number of miRNAs and 248 is the number of diseases. The more details are shown in Fig. 2.

**Fig. 2.**
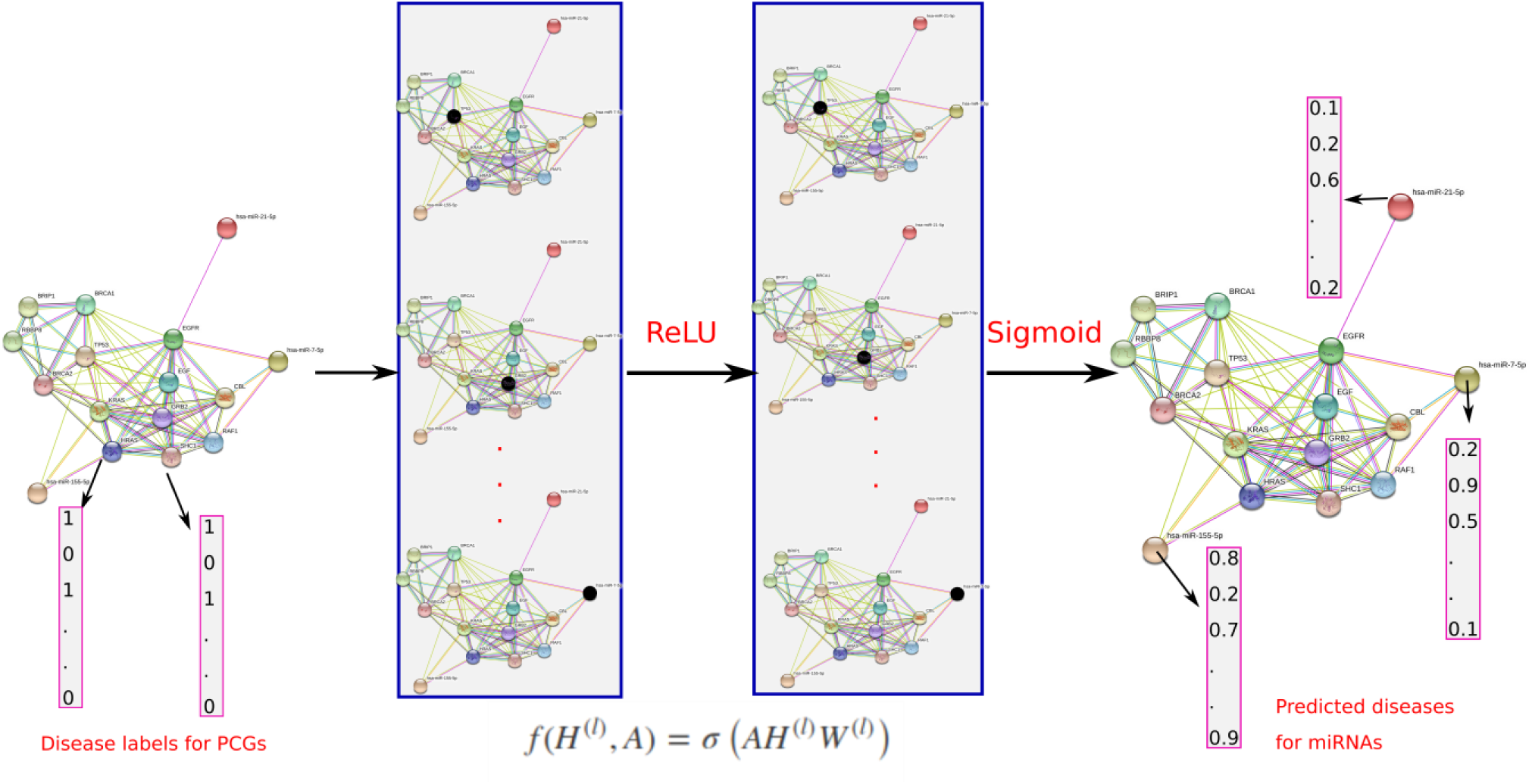
The flowchart of DimiG with 2-layers GCN. When doing forward propagation, the embedding of the dark node in each network is the weighted sum of the embedding of its neighbors, where all nodes in the network are updated simultaneously. The label is multi-hot vector indicating the presence of diseases. In the end, DimiG can infer the probability between diseases and unlabeled miRNAs. The embedded network example is from STRING database (Szklarczyk, et al., 2015).

### 2.6 Baseline methods

We formulate disease-miRNA association prediction as node classification in a graph, where nodes of PCGs are assigned multiple diseases as labels, and nodes of miRNAs are used as the test set. The miRNA-disease associations are completely not involved in model training of DimiG. To demonstrate the power of DimiG, we compare with two existing similar baseline methods that do not require disease information for miRNAs, and two variants of our DimiG model:

1. miRPD: it scores miRNA–disease associations by network analysis of miRNA–protein associations and protein–disease associations from text mining (Mork, et al., 2014). We downloaded the inferred associations between diseases and miRNAs from their websites. In this study, we downloaded full dataset including croft_full.tsv.gz, miranda_full.tsv.gz and targetscan_full.tsv.gz. The three sets give the association scores between diseases and miRNAs, they are called miRBD-C, miRBD-M and miRBD-T, respectively.
2. CNC: it is a co-expression-based method under guilt-by-association framework. For each disease *d* and miRNA, CNC consists of the following steps: 1) Calculate Pearson’s correlation coefficients (PCC) between D-associated PCGs and the miRNA. 2) Keep only those co-expression PCGs with absolute PCC > 0.3 and *P*-value < 0.01 (Liu and Zhao, 2016). 3) Calculate the mean value of up to K largest PCC value as the score for the miRNA with disease *d*.
3. Two variant methods of DimiG: instead of using expression profiles across tissues as node features, we used one-hot encoding of all nodes. Simply, we use the identity matrix as the node feature matrix *X*. This variant method is called DimiG-I. In addition, we also construct another variant method DimiG-C, which combines the expression profiles and one-hot encoding as node features.

After obtaining the associated scores between diseases and miRNAs, we extracted scores for those positive and negative disease-miRNA pairs in the independent test set, and these scores are pooled together to calculate the area under the receiver operating characteristic curve (AUC) and the area under precision-recall curve (AUPRC). For miRPD, only those disease-miRNA pairs with scores in the download files are kept.

We also compare DimiG with other two state-of-the-art supervised methods BNPMDA (Chen, et al., 2018) and DRMDA, both use miRNA-disease associations for model training. BNPMDA trains a bipartite network recommendation method on known disease-miRNA associations by integrating miRNA and disease similarity. DRMDA further uses stacked autoencoder to learn deep representation for diseases and miRNAs, which are fed into a support vector machine for predicting disease-miRNA associations. Both methods are trained on disease-miRNA associations from the database HMDD v2.0.

As mentioned above, cross-validation may yield biased performance. Thus, we evaluate how BNPMDA and DRMDA generalize to unseen miRNA-disease associations in the training set. We download the prediction association scores for available miRNAs and diseases from the supplementary material of these two studies, and the disease names are mapped to the disease ontology ID. Of the 1034 miRNAs and 248 diseases in this study, 304 miRNAs and 55 diseases are in the score matrix predicted by BNPMDA and DRMDA. Instead, the DimiG does not use any disease-miRNA association information for model training, to make a fair and objective comparison, we evaluate the prediction performance of DimiG, BNPMDA and DRMDA on a dataset consisting of newly recorded verified miRNA-disease associations in HDMM v3.0 but not in HMDD v2.0. We call this dataset as an unseen disease-miRNA set, which is freely available at https://github.com/xypan1232/DimiG/tree/master/data. In this data set, there are 1954 disease-miRNA pairs consisting of 977 positives and 977 negatives, and it is also balanced not only overall but also for each disease. It should be noted that this data set may still have some possible dependent associations with the training set derived from HMDD v2.0.

### 2.7 Experimental settings

In this study, we modify the implemented GCN from https://github.com/tkipf/pygcn to support multi-label classification using PyTorch framework. DimiG consists of two-layer GCNs. The last layer is the sigmoid layer with number of neurons 248, which is the number of diseases as labels. We use Adam to optimize the Binary Cross Entropy between the targets (Diederik and Jimmy, 2015). We use grid search to select the best parameters for learning rate, weight_decay and dropout probability.

Of the 6,188 PCGs, we keep 80% of PCGs as the training set, 20% of PCGs as the validation set and all 1034 miRNAs as the test set, which is used to evaluate the performance of DimiG for predicting disease-associated miRNAs. For each PCG, the label is a 248-dimensional multi-hot vector indicating the presence of diseases associated with this PCG.

### 2.8 Case studies

In this study, we further evaluate the DimiG for predicting associated miRNAs for three types of diseases, i.e., prostate cancer, lung cancer and inflammatory bowel disease (IBD). For prostate cancer, of all the 1034 miRNAs, 176 miRNAs are associated with prostate cancer in HMDD v3.0, which are used as the positives. For negative control, we randomly select other 176 miRNAs not associated with prostate cancer in the HMDD v3.0 as negatives. The 176 positives and 176 negatives comprise the prostate cancer specific dataset. Then we pool the predicted scores by DimiG of these total 352 miRNAs for prostate cancer and calculate the AUC scores for performance evaluation.

We also use the same way to construct lung cancer and IBD specific datasets. Of the 1034 miRNAs, 172 and 18 are recorded as association with lung cancer and IBD in HMDD v3.0 as positives, respectively. We also randomly selected the same number of miRNAs as negatives. For each of the three diseases, we will also check the overlap between the top 50 predicted miRNAs and its associated miRNAs in HMDD v3.0.

In addition to the HMDD database, we also extract verified disease-miRNA associations from cancer database dbDEMC v2.0 (Yang, et al., 2017). We download disease-miRNA associations derived from low-throughput methods. Of the 1034 miRNAs, 15 and 16 miRNAs are recorded as association with prostate cancer and lung cancer in dbDEMC v2.0, respectively. These two numbers are much fewer than that of HMDD v3.0, it is because only disease-miRNA associations verified by low-throughput methods before 2017 are used for dbDEMC v2.0. Similarly, for each disease, we also randomly selected the same number of miRNAs as negative controls.

## 3 Results

In this study, we first evaluate the prediction performance of DimiG on four tissue expression data with different number of tissues. Then we compare DimiG with other baseline methods that do not use disease-miRNA associations for model training on the independent test set, and further compared to BNPMDA and DRMDA trained using disease-miRNA associations on the unseen disease-miRNA set. Lastly, we present three case studies for prostate cancer, lung cancer and IBD.

### 3.1 The performance of DimiG on four tissue expression datasets

We first checked the impact of number of epochs on training DimiG. As shown in Fig. 3A, the training and validation loss converge to the same as the number of epochs approaches to 50. Thus, in this study, we use epochs 50 for our below experiments. It should be noted that the training loss is larger than validation loss during first 30 epochs, which is also observed for the citation network data in the GCN paper (Kipf and Welling, 2017). One possible reason is regularization (e.g. dropout), which is used during training but not during validation. Another potential reason is that the features of the genes have certain discriminative power, and they behave similarly to other genes in the training set. After several epochs, the learned node features are propagated well from interacting neighbor nodes for both training and validation genes.

**Fig. 3.**
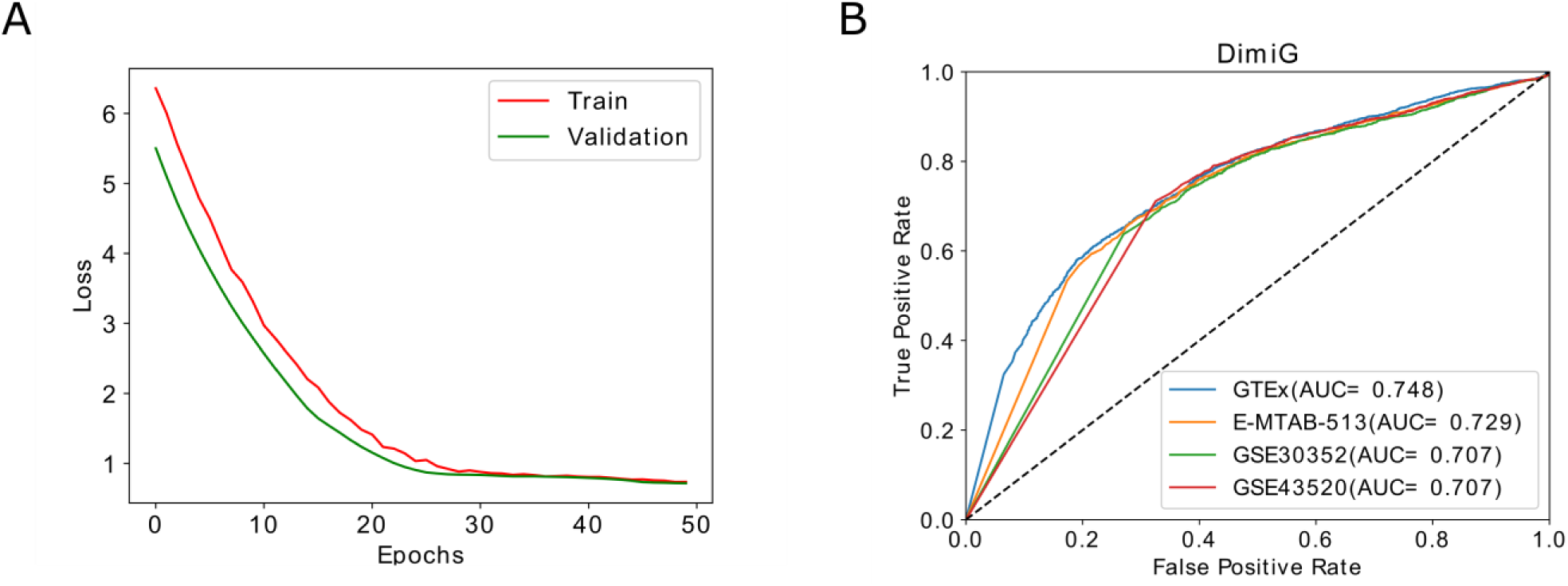
The training and performance of DimiG. A) The training and validation loss change with the number of epochs on GTEx dataset. B) The ROC curve of DimiG using expression profiles from different dataset, where GTEx, E-MTAB-513, GSE30352 and GSE43520 covers 53, 16, 6 and 4 human tissues, respectively.

Grid search approach is used to select the best parameters for DimiG on validation set, where we search the learning rate with values of [0.001, 0.005, 0.0001, 0.0005], number of neurons in hidden layer with values of [248, 496, 744, 992, 1984], weight_decay with values of [0.001, 0.005, 0.0001] and Dropout with values of [0.5, 0.7, 0.8]. As shown in Supplementary Fig. S2, we yield the best AUC when learning rate=0.0001, number of neurons = 744, weight_decay = 0.005 and Dropout = 0.8, which are finally used in our DimiG model.

As shown Fig. 3B, DimiG yields the AUCs of 0.748, 0.729, 0.707 and 0.707 on GTEx, E-MTAB-513, GSE30352 and GSE43520, respectively. It yields the best performance on GTEx, which covers 53 tissues and thus contains more complete tissues. Compared to GTEx, DimiG achieves lower performance on GSE43520. It is presumably because that GSE43520 only covers four tissues, many tissues associated with certain diseases are missing.

When expression profiles from fewer tissues represent node features, DimiG easily suffers from noises that may be caused by sequencing errors. Of more interest, the area of the ROC curve with low false positive rate for GTEx is much bigger than other three datasets, this region is especially important for evaluating predictive models. The results indicate that expression profiles across more tissues as node features can improve the prediction performance for disease-miRNA associations.

### 3.2 Comparing DimiG with other baseline methods that do not use disease-miRNA associations for model training

In DimiG, tissue expression profiles are used as node features. Note its variant of DimiG-I just uses the one-hot encoding as the node features. As shown in Fig. 4A, the proposed final DimiG yields an AUC of 0.748, which is an increase of 6% over AUC 0.706 of DimiG-I. These results demonstrate that expression profiles across tissues are very informative for inferring disease-miRNA associations. Another variant DimiG-C combines the expression profiles and one-hot encoding of nodes as node features and yields an AUC of 0.728, which is better than DimiG-I, but still is worse than DimiG. These results demonstrate that simply concatenating the expression profile and one-hot encoding of nodes not only introduces computational burden, but may also decrease the prediction performance. One possible reason is that the one-hot encoding of nodes has no correlations with the expression profiles, making the GCN cannot encode the node features well.

**Fig. 4.**
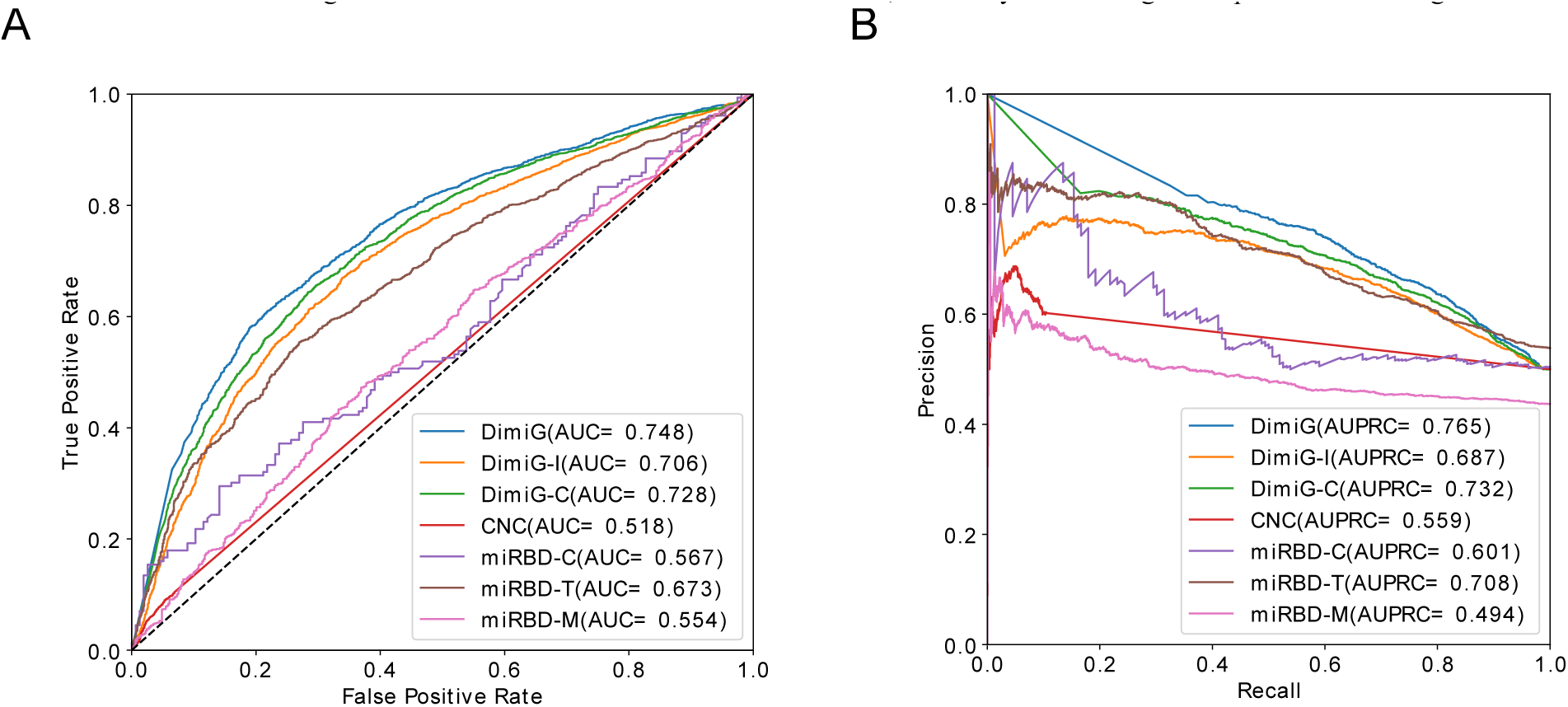
The performance of DimiG using expression profiles as node features from GTEx and baseline methods. A) ROC curve. B) Precision-recall curve. AUC is the area under ROC curve and AUPRC is the area under precision-recall curve.

We also compare DimiG with another published method miRPD, which provides association scores between diseases and miRNAs derived from three different sources of miRNA-PCGs. As shown in Fig. 4A, DimiG outperforms the best AUC 0.673 of miRPD-T by 11.1%. Of the three miRPD-based methods, miRPD -T yields the best AUC 0.673, and it is based on the miRNA-PCGs predicted by TargetScan. The miRPD-C and miRPD-M yield similar AUC, which is much worse than miRPD-T. One potential reason is that miRPD-C infers association scores only based on a small number of verified miRNA-PCG. Although miRPD-M uses similar number of miRNA-PCG, but it may suffer to high false positives than TargetScan.

In addition, our results show that the coexpression-based CNC method yields the performance AUC=0.518, suggesting the co-expressed PCGs with miRNAs are not enough for identifying high-confidence disease-miRNA associations. It is because CNC is only based on expression profiles, interactions information of miRNAs is completely ignored.

We also calculate the AUPRC of different methods. As shown in Fig. 4B, DimiG yields the best AUPRC of 0.765, which is also better than its two variants DimiG-I and DimiG-C. In addition, compared to state-of-the-art method miPRD, the best miRBD-T yields the AUPRC of 0.708, which is ∼ 9% worse than AUPRC 0.765 of DimiG.

All the above results show that DimiG is able to achieve better performance for inferring disease-miRNA associations, which is not a surprise since that GCNs can better integrate interaction network data and tissue expression profiles, and the GCNs can operate on graphs similarly to CNNs on images, and takes the features and connectivity of nearby nodes into account.

### 3.3 Performance comparison of DimiG on the unseen disease-miRNA set

We also compare DimiG with other two state-of-the-art supervised methods BNPMDA and DRMDA on the unseen disease-miRNA set, both BNPMDA and DRMDA use disease-miRNA associations for model training. As shown in Fig. 5, on this unseen disease-miRNA set, DimiG yields an AUC of 0.710 and an AUPRC of 0.724, which are better than the performance of BNPMDA with an AUC 0.686 and an AUPRC of 0.698, and the performance of DRMDA with an AUC of 0.708 and an AUC of 0.715. The results indicate that DimiG outperforms the state-of-the-art supervised methods on inferring new disease-miRNA associations. The AUC 0.687 of BNPMDA on this unseen disease-miRNA set is lower than the reported 5-fold cross-validation AUC of 0.898. Similarly, the AUC 0.708 of DRMDA is also lower than the reported 5-fold cross-validation AUC of 0.916. The results demonstrate that the methods trained on disease-miRNA associations may yield biased cross-validation performance, which could not reflect the methods’ actual ability to predict unseen miRNA-disease associations and generalize well to new miRNA-disease associations, as observed in (Park and Marcotte, 2012). Since DimiG does not use any disease-miRNA associations, the performance of DimiG (an AUC of 0.710) on this unseen disease-miRNA set is consistent with the performance (an AUC of 0.748) on the full independent test set constructed from HMDD v3.0. The results indicate the reported performance reflects DimiG’s ability to infer new disease-miRNA associations.

**Fig. 5.**
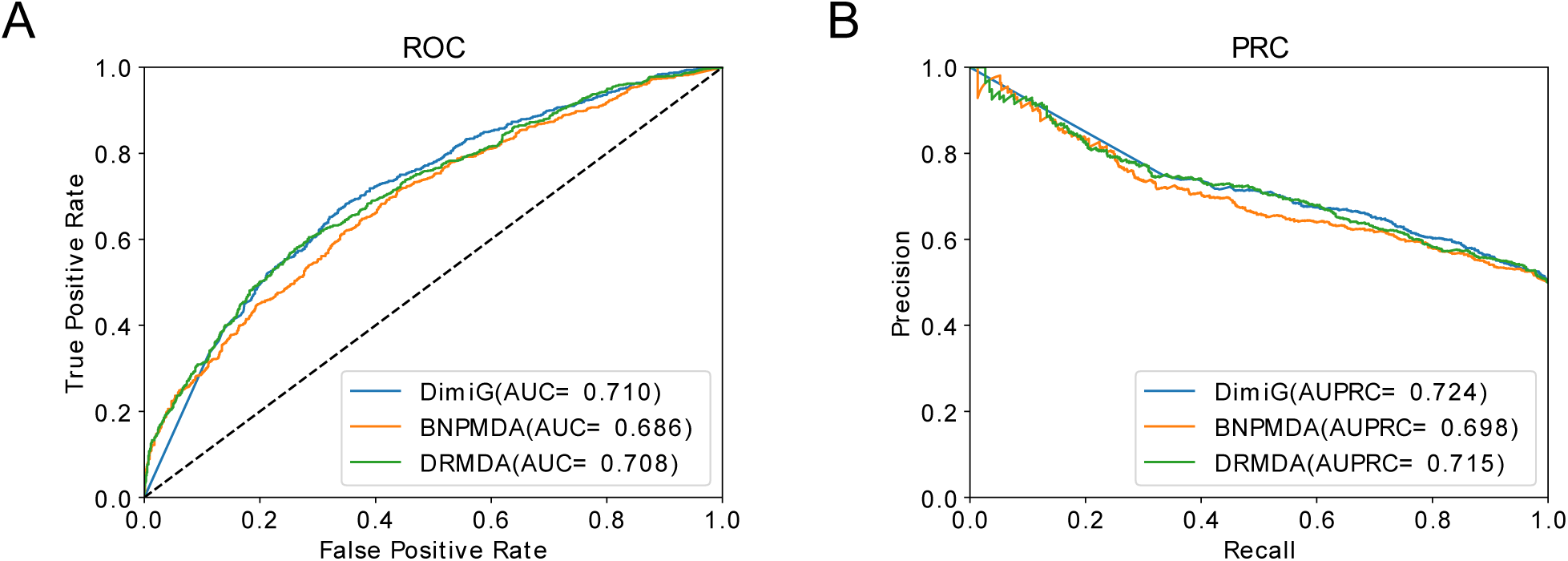
The performance of DimiG, BNPMDA and DRMDA on the unseen disease-miRNA dataset. Here, DimiG uses expression profiles as node features from GTEx, BNPMDA and DRMDA are trained on known disease-miRNA associations. A) ROC curve. B) Precision-recall curve.

### 3.4 Case studies

We present three case studies of miRNAs associated with prostate cancer, lung cancer and IBD to demonstrate the applicability of DimiG for inferring disease-associated miRNAs. The top predicted candidates for these three diseases are checked with verified associations from literature and public databases, HMDD v3.0 and dbDEMC v2.0. In addition, we evaluate the disease-specific prediction performance of DimiG.

#### 3.4.1 Prostate cancer

We first investigate the prediction of prostate cancer associated miRNAs from DimiG. We list the top 10 miRNA candidates predicted by DimiG in Table 1, which are all supported by the literature or database dbDEMC v2.0. For example, a meta-analysis shows that the top 1st miR-939 and top 8^th^ miR-661 are downregulated, and the top 9^th^ miR-637 is upregulated in recurrent prostate cancer (Pashaei, et al., 2017). This study also finds prostate cancer associated CTNNB1 (Anastas and Moon, 2013) is one hub gene for their interacting targets in a gene network. That’s to say, the predicted microRNA genes by DimiG of Table 1 are well-consistent with existing knowledge.

**Table 1.** The top 10 candidate prostate cancer related miRNAs predicted by DimiG and their support evidences in literature.

| Rank | MiRNA name | Reason | Support evidence (PMID ID) |
| --- | --- | --- | --- |
| 1 | miR-939 | Downregulated | 28651018 |
| 2 | miR-93 | Overexpressed in prostate cancer, and downregulate capicua levels to promote prostate cancer progression | 26124181 |
| 3 | miR-92a | Regulate tumor suppressor PTEN | 24098737 |
| 4 | miR-874 | Upregulated | dbDEMC v2.0 |
| 5 | miR-766 | Downregulated | dbDEMC v2.0 |
| 6 | miR-765 | Mimic the expression of oncogenic HMGA1 in prostate cancer | 24837491 |
| 7 | miR-744 | Promote prostate cancer progression by activating Wnt/β-catenin pathway | 28107193 |
| 8 | miR-661 | Downregulated | 28651018 |
| 9 | miR-637 | Upregulated | 28651018 |
| 10 | miR-625 | Upregulated | dbDEMC v2.0 |

In another study, the top 2rd miRNA miR-93 is frequently overexpressed in prostate cancer and down-regulates capicua levels (Choi, et al., 2015). In dbDEMC v2.0, three miRNAs miR-874, miR-766 and miR-625 are differentially expressed in prostate cancer. Of them, miR-625 and miR-874 share the same gene target HMGA1 in prediction channel of RAIN database with miR-765. The remaining three miRNAs in the top 10 candidates either regulate prostate cancer associated genes or activate prostate cancer associated pathway. We have also note that of the top 10 miRNAs, only miR-92a and miR-765 are recorded in HMDD v3.0, and the others are not. These results indicate that DimiG can infer novel disease-miRNA associations currently not in the curated databases.

As shown in Fig. 6A, of the 176 prostate cancer associated miRNAs in HMDD v3.0, 21 are in the top 50 predicted candidates by DimiG. Six of the 15 prostate cancer associated miRNAs in dbDEMC v2.0 belong to the top 50 candidates predicted by DimiG (Supplementary Fig. S3A). On the prostate cancer specific dataset constructed from HMDD v3.0, DimiG yields an AUC of 0.724, 0.697, 0.675 and 0.664 on GTEx, E-MTAB-513, GSE30352 and GSE43520, respectively (Fig. 6B). Similarly, on the dataset constructed from dbDEMC v2.0, DimiG achieves an AUC of 0.844, 0.836, 0.729 and 0.684 on GTEx, E-MTAB-513, GSE30352 and GSE43520 for prostate cancer, respectively (Supplementary Fig. S3B). The performance is better than that on the dataset constructed from HMDD v3.0 since the extracted disease-miRNA associations are experimentally verified using low-throughput methods and more reliable. We can observe that more tissues can provide informative clues for predicting prostate cancer associated miRNAs, even some tissues maybe consider not relevant to prostate cancer. The results further demonstrate the power of DimiG.

**Fig. 6.**
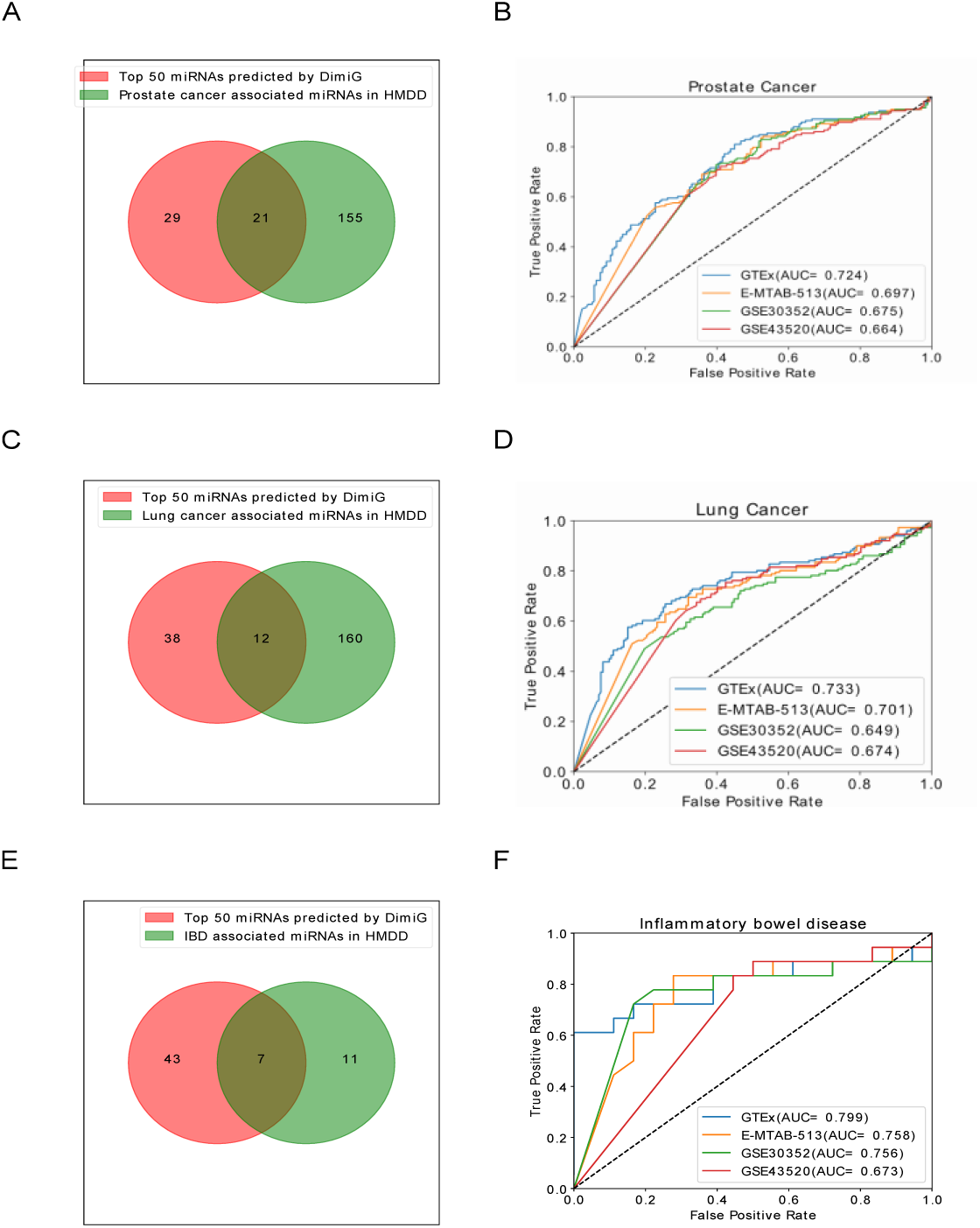
Venn diagram and ROC curve for predicting associated miRNAs for prostate cancer, lung cancer and IBD using DimiG. A) The overlap between the top 50 predicted miRNAs by DimiG and prostate cancer associated miRNAs in HMDD v3.0. B) ROC curve for predicting prostate cancer associated miRNAs using four tissue expression datasets. C) and D) Venn diagram and ROC for Lung cancer, respectively. E) and F) Venn diagram and ROC for Inflammatory bowel disease (IBD), respectively.

#### 3.4.2 Lung cancer

We investigate the prediction ability of DimiG for lung cancer associated miRNAs. There are 172 such miRNAs recorded in the HMDD v3.0 for lung cancer. As shown in Fig. 6C, of the 172 miRNAs, 12 miRNAs are in the top 50 lung cancer associated miRNAs predicted by DimiG. Then we evaluate the prediction performance on the dataset consisting of 172 lung cancer associated miRNAs and 172 miRNAs not associated with lung cancer in HMDD v3.0. As shown in Fig. 6D, DimiG yields an AUC of 0.733, 0.701, 0.649 and 0.674 for GTEx, E-MTAB-513, GSE30352 and GSE43520, respectively.

We further report the prediction ability on another lung cancer-specific dataset derived from dbDEMC v2.0. In this dataset, there are 16 miRNAs associated with lung cancer, and other 16 miRNAs not associated with lung cancer in dbDEMC 2.0. Of the 16 miRNAs associated with lung cancer, one is in the top 50 predicted miRNAs by DimiG (Supplementary Fig. S3C). As shown in Supplementary Fig. S3D, DimiG achieves an AUC of 0.925, 0.808, 0.806 and 0.869 for GTEx, E-MTAB-513, GSE30352 and GSE43520, respectively. The results also indicate informative clues can be captured in more tissues for predicting lung cancer associated miRNAs.

#### 3.4.3 Inflammatory bowel disease

Beside the cancer disease, we also investigate DimiG’s prediction ability for another non-cancer disease IBD. In this study, we predicted associated diseases for 1034 miRNAs, of which 18 miRNAs are deposited for IBD. Seven of the 18 miRNAs are in the top 50 miRNAs predicted by DimiG (Fig. 6E). As shown in Fig. 6F, DimiG yields an AUC of 0.799, 0.758, 0.756 and 0.673 for GTEx, E-MTAB-513, GSE30352 and GSE43520, respectively. The results show that we can use DimiG for other non-cancer diseases.

## 4 Discussion

In this study, we present a semi-supervised multi-label GCN framework to integrate heterogeneous networks of tissue expression profiles, miRNA-PCG interactions, PCG-PCG interactions, disease-PCG interaction to infer disease-associated miRNAs. The whole pipeline is under the context of interaction network, where disease-miRNA associations are completely not involved in model training. CNN cannot directly process the non-Euclidean domain data, like network data. But GCN can handle these types of data, they are specially designed to extract abstract features from network data. We demonstrate that cross-validating the methods trained on disease-miRNA associations yields an overestimated performance. Our results demonstrate that DimiG outperforms other state-of-the-art methods, which does not require disease-miRNA associations information, and two methods trained on disease-miRNA associations.

In this study, we need set several cut-off values (Fig. 1) for PCG-PCG interactions from STRING database, PCG-miRNA interactions from RAIN database and disease-PCG associations from DISEASES database, respectively. All the three databases are developed by the same group, they use the similar data quality control, and the confidence values are all scored using the similar pipeline from multiple channels, including experiments, knowledge, text mining and prediction. Thus the constructed graph should follow the GCN’s assumption that it is a simple one-modal graph, in which all nodes are of the same type (all nodes are genes), and all edges have the same semantic meaning (Kipf and Welling, 2017).

As shown in Supplementary Fig. S4, for the low cutoff values of STRING with 300 and RAIN with 0.10, DimiG yields much lower performance with AUC 0.653, it is because lower cutoff may introduce more false positive interactions. For cutoff value of STRING greater than 400 and cutoff value of RAIN greater than 0.15, DimiG yields similar AUCs. DimiG yields the best AUC 0.754 at cutoff values 400 and 0.20 for STRING and RAIN, respectively, but these higher cutoff values will lead to fewer miRNAs. Thus, we use cutoff value 400 of STRING and 0.15 of RAIN in this study, and DimiG yields an AUC 0f 0.748 and is capable of finding more miRNAs, which is just a little lower than 0.753 with cutoff value 400 and 0.20 for STRING and RAIN databases as the tradeoff. We also evaluate the impact of confidence thresholds 1.5 and 2.5 for disease-PCG associations on the performance of DimiG. As shown in Supplementary Fig. S5, DimiG yields an AUC of 0.719 and 0.742 for threshold 1.5 and 2.5, respectively, both are lower than 0.748 when using threshold 2. The possible reason is that lower confidence threshold 1.5 introduces more false positives for model training, and higher threshold 2.5 makes the number of training samples much fewer, which are both not good for training machine learning model.

DimiG does not require disease-miRNA associations for model training, but it requires the miRNA-PCG interactions to construct the graph. Each miRNA must have at least one interacting PCG, all nodes, including miRNAs and PCGs, need be present during the training. Thus, the trained node embedding can be propagated to miRNAs and further used for inferring miRNA-associated diseases. This precondition makes us to discard some miRNAs, and DimiG cannot infer associated diseases for these miRNAs without interacting PCGs. In addition, more and more novel miRNAs are being discovered, and their interactions with PCGs may not be readily available. Thus, the trained models cannot be generalized to miRNAs not in the graph. Luckily, some recent GCN models have tried to solve this issue, which are trained on a set of nodes and generalized to any augmentation of the graph (Hamilton, et al., 2017). This inductive GCN model can be applied for novel miRNAs not in the graph in our future study.

DimiG does not use miRNA-disease associations during model training. There are other computational models with reported cross-validation AUC over 0.8, in which disease-miRNA associations are involved in model training. We demonstrate that cross-validation could report overoptimistic performance of the methods and could not generalize well to unseen disease-miRNA associations. Disease-miRNA associations can be incorporated into model training, however, randomly dividing the disease-miRNA pairs into the training and test sets for cross-validation could be biased. To better evaluate the performance of one model, a strictly independent test set should be at least constructed, e.g. the model is trained on all data published up to a specific year, and predictions are evaluated on data published after that, or the model is trained on data in the older version of database and evaluated on those new added disease-miRNA associations in the updated database. In addition, DimiG predicts associated miRNAs for 248 diseases in one model. Many previous methods formulate the disease-miRNA prediction as binary classification problems, and they require constructing negative disease-miRNA associations, which may introduce false negatives into model training.

In this study, we used only expression profiles across tissues as node features. Some studies have revealed that functional domain information can assist identifying disease-associated miRNAs (Yang, et al., 2018). In future work, we can combine the gene ontology (GO) information and expression values across tissues into the node features, or ensemble the two GCN models trained on each representation, which is expected to further improve the prediction performance of DimiG.

## 5 Conclusion

In this study, we present a semi-supervised multi-label learning framework DimiG to integrate interaction data for inferring miRNA-associated diseases. DimiG does not use any disease-miRNA associations for model training. This new approach achieves promising performance and outperforms other baseline methods not trained on disease-miRNA associations with a large margin on our benchmark dataset. We also observe that cross-validation performance of methods trained on known disease-miRNA associations could be overestimated and could not reflect their actual abilities for inferring new disease-miRNA associations. Our results demonstrate that the tissue expression profiles can provide informative signals for inferring disease-miRNA associations. We expect DimiG to be used to discover novel miRNA biomarkers for diseases and the framework can be extended to other tasks based on network data, e.g. functional annotations of proteins.

## Funding

This work was supported by the National Key Research and Development Program of China (No. 2018YFC0910500), the National Natural Science Foundation of China (No. 61725302, 61671288, 91530321, 61603161), the Science and Technology Commission of Shanghai Municipality (No. 16ZR1448700, 16JC1404300, 17JC1403500).

## Conflict of Interest

none declared.

